# How emotional memory affects face remembering: an ERP investigation

**DOI:** 10.1101/208256

**Authors:** Alice Mado Proverbio, Maria Elide Vanutelli, Simone Viganò

## Abstract

It is known that the longer an information has been memorized, the stronger is its memory trace. At the same time it is known that the emotionally-valenced information has a stronger memory trace than neutral one. Interactive effects between these factors are largely unknown. In this study electrical neuroimaging signals were recorded in healthy controls to explore the neural mechanism of memory for faces of different emotional valence, sex, learning context and temporal recency. In a study phase participants familiarized with the fictional police dossier of 10 victims of dramatic deaths including homicide or earthquake (depicted as attractive and nice persons of about 45 years), twice a day for seven days before EEG recordings. Hundreds of famous movie stars (overlearned), victims (recent) and new faces were presented in an old/new recognition task. ERP responses showed a modulation of anterior N2 and FN400 as a function of face familiarity (with no differences between overlearned and recent faces), while parietal late positivity was sensitive to engram temporal recency (it was much larger to overlearned than recent faces, paralleling behavioral data). However, LP amplitude did not differ to overlearned vs. recent same-sex faces, showing how emotional memory can enhance faces remembering. A late frontal negativity revealed sensitive to source memory.

## Introduction

The goal of present study was to compare the time course and functional properties of electrophysiological responses elicited during encoding and retrieval of faces in a paradigm in which memory for old items (such as the faces of popular movie stars, known for at least 5-8 consecutive years), was compared with that of recently acquired information (faces of fictional characters studied during the week preceding the EEG recording) or novel information (faces of previously unknown people). In this way the effect of temporal recency of faces on remembering was investigated, by considering the gradient: overlearned, recent, and new. At this regard, previous literature (e.g., Moreton and Ward, 2010) investigated the extent to which retrieval from long-term memory differs as a function of the retention time. In Moreton’s and Ward’s study (2010), for example, participants were given 4 minutes to recall autobiographical events from the last 5 weeks, the last 5 months, or the last 5 years, and the results showed similar retrieval rates. This results has been interpreted in the light of the so-called SIMPLE model (Brown et al., 2007), according to which correct memory recognitions would be a function of the relative temporal distinctiveness of the items to be remembered, not just of their recency. However, the neuroscientific literature provides evidence of a different neural representation of short vs. long term memories. For example Bergmann et al. (2012), in an event-related fMRI study, found that right anterior hippocampus and left inferior frontal activations predicted successful long term memory for pairs of faces and houses, while activations of the left parahippocampal gyrus including the fusiform gyrus predicted subsequent accuracy on working memory decisions. Furthermore, it has long been known that in case of damage to hippocampal or medial temporal regions, the retrograde amnesia deficit involves only the recent past (for example memories relative to the last few years preceding the cerebral insult) and not the remote past of the patient, thus indicating a greater strength of older memory traces (“Consolidation theory” by Manns et al., 2003, see also Markowitsch and Staniloiu, 2013; Kopelman and Bright, 2012). In addition, according to the “Episodic-to-semantic shift” theory, also called “semanticisation”, episodic memories would acquire a more ‘semantic’ form as they get older, thereby protecting them against the effects of brain damage (Cermak, 1984) or oblivion. Furthermore neurophysiological data have provided multiple evidences that brain encoding of very recent memories would have a bioelectrical nature (e.g. Zamora et al., 2016), while the deep encoding in LTM would be based on plastic structural changes and dendritic formations, also called long-term potentiation (LTP).

Notwithstanding that, not many studies in humans have directly compared the neural markers of remembering information of different temporal recency, especially with the ERPs (event-related potentials) technique, which is actually very sensitive to both recollection and familiarity processes.

EEG and neuroimaging studies have suggested the existence of at least two independent processes underlying recognition memory, namely recollection and familiarity (Henson et al., 1999). Recollection reflects the conscious retrieval of contextual information about a specific episode, while familiarity would refer to an acontextual feeling of knowing. Both processes are thought to be related to two spatiotemporally different ERP effects (FN400 and LP), namely the early mid-frontal negativity (familiarity) and the late parietal old/new effect (recollection) (Evans and Wilding, 2012; Rugg et al., 1996; Wilding & Rugg, 1996; Wilding et al., 2003; Hoppstädter et al., 2015). According to the dual-process model of recognition memory, judgments produced on the basis of familiarity are quick, automatic, and reflect a general feeling of knowing that a stimulus was previously presented. On the other hand, recollection requires actively reporting details about studied items and results in slow, more effortful judgments (Aggleton and Brown, 1999; Atkinson and Juola, 1974; Yonelinas et al., 1999, 2002). Some authors have argued that F/N400 response does not necessarily reflect only familiarity but also conceptual priming, which is an implicit-memory phenomenon that often occurs together with familiarity (e.g., Paller et al., 2007; Wang and Yonelinas, 2012; Jäger et al., 2006). For example Voss and Paller (2006) investigated ERP correlates of memory for famous faces with a paradigm in which, during learning, some set of faces were presented along with corresponding biographical information. Memory was therefore enhanced via conceptual priming so that old/new recognitions were faster and more accurate for primed compared with unprimed famous faces. Interestingly F/N400 mid frontal effect reflected this manipulation. In this venue F/N400 might reflect a type of source memory (being it sensitive to learning circumstances.)

The present research applied the ERPs technique to explore the temporal course of brain bioelectrical activity during recognition of faces of different emotional valence, sex, learning context and temporal recency. The study involved the presentation of 300 faces: some belonged to famous people (movie and TV stars), others were learned during a study phase starting 1 week before the experiment (very recent material), and the remaining were new faces (no familiarity). Faces to be studied (previously unknown) were presented as victims of murders or accidents with a description provided in the form of a police dossier featuring a short text accompanied by several pictures of the victims. This experimental manipulation was included to explore the effect of emotional memory in the episodic encoding of faces. Indeed, convergent evidence suggests that emotional events attain a privileged status in memory (LaBar & Cabeza, 2006) and that memory tends to be better for emotionally arousing information than for neutral information (Buchanan & Lovallo, 2001).

Recently, Mueller and Pizzagalli (2016) have provided direct electrophysiological evidence (via intracranial recordings) that faces that had been fear-conditioned (to become threatening stimuli) evoke increased activity in or in proximity of the left fusiform gyrus as early as 80 ms post-stimulus after presentation. The effect is visible starting from the same day of conditioning and, remarkably, it is still very strong one year after the fear-conditioning. In another experiment Proverbio et al. (2016) investigated the effect of social prejudice on memory for faces, by presenting hundreds of faces of males and females in association with a derogatory or a positive description, and found that faces associated with a negative (vs. positive) prejudice elicited larger anterior negativities during encoding, smaller FN400s and larger old/new parietal effects during recollection. The larger potentials for negatively-valenced faces were associated to increased activations of the limbic and parahippocampal areas. Indeed other variables, besides temporal recency of learning, seem to affect face remembering. For example, it has been demonstrated that face recognition can be influenced by the context present at the time of encoding, like the lightning (see a discussion in Megreya & Burton, 2008), the type of the facial expression (D’Argembeau & Van der Linden, 2011), the level of the observer empathy (Bate et al., 2009) or face gender (Proverbio et al., 2010).

According to previous ERP literature, we predicted a significant modulation of FN400 and LPC components (old/new recognition effects) and were interested in investigating the impact of emotional valence and recency on the ability to remember faces, in the hypothesis that emotionally valenced (but recently acquired) faces might be remembered equally well than robustly coded information (e.g., extremely popular faces). In detail, a frontally distributed N400 (F/N400), has been directly related to the concept of familiarity. For example, Curran (Curran, 1999; 2000) found an enhanced negative response between 300-500 ms post-stimulus at scalp anterior regions, which was larger to new than old items. Additionally, we expected a modulation of an earlier familiarity-related response (frontal N2) recently found to precede F/N400 effects, and being also larger to unknown than familiar items (Lawson et al., 2012).

Additionally we also wished to investigate whether face sex significantly affected memory recollection. Indeed previous research on emotions and empathic resonance mechanisms showed that some specific factors can facilitate emotional mirroring, like perceived similarity (e.g., sex of viewer); it would influence empathic responding through the tendency to identify more closely with others who appear as being more similar in features such as personality (Gruen and Mendelsohn, 1986) or appearance (Bufalari et al., 2015). This effect might emerge in the form of a face sex effect in the present study, by considering the own/different variable with respect to the viewer. Indeed, according to the neuroimaging literature self-referential information would be represented in distinct brain loci (as compared to social information about other people) and would be better remembered. The self-reference effect (SRE), described as the better performance in recalling or recognizing memorized materials when it refers to the self, has been explored with neuroimaging techniques, which have identified a ventral-to-dorsal gradient within the medial prefrontal cortex, for representing self vs. other information (Sul et al., 2015). On the basis of this literature we might expect a better recall of same-sex faces, sharing sexual gender with the viewer.

## Materials and methods

### Participants

Twenty-two university students (11 males and 11 females) volunteered for this experiment. Females ranged in age from 20 to 26 years (mean age=22.91 years, SD=1.76) and had a high level of education (15.73 years in school, SD=1.01). Males ranged in age from 20 to 27 years (mean age=24.18 years, SD=2.05) with the same level of education as females (15.82 years in school, SD=1.66). All participants had heterosexual preferences as ascertained by a written questionnaire, had normal or corrected-to-normal vision with right eye dominance, and were right-handed as assessed by the Edinburgh Inventory with no left-handed relatives. Experiments were conducted with the understanding and written consent of each participant according to the Declaration of Helsinki (BMJ 1991; 302: 1194) with approval from the Ethical Committee of University of Milano-Bicocca and in compliance with APA ethical standards for the treatment of human volunteers (1992, American Psychological Association). All participants received academic credits for their participation. Data from 3 male subjects were subsequently discarded because of excessive eye movements and electroencephalogram (EEG) artifacts (more than 20% of trials rejected). The final sample included nineteen university students (8 males and 11 females). Males ranged in age from 20 to 27 years (mean age=24 years, SD=2.27) with a mean of 15.9 years in school (SD=1.96).

### Stimuli

Stimulus set included three different face categories: famous (overlearned), recently studied and new faces. Famous faces belonged to popular actors or TV anchors who have been extremely popular for at least 5 years (namely within the 2005-2010 period) but still very popular for people of about 23 years of age (in 2013). For example, when considering actors, it was verified the date of hit movies or TV shows that put them in the spotlight, (for example the famous TV series “Doctor House” played by Hugh Laurie (broadcasted from 2005 to 2012 in Italy), or the movie “Speed” (2006) played by Sandra Bullock, or the movie “The Devil wears Prada” (2006) played by Meryl Streep. New faces, instead, were completely unknown to the participants and were downloaded from free online databases. Recently studies faces (previously unknown) were memorized by participants during a study phase lasting 1 week and preceding the experimental session, in which they were depicted as victims of a homicide or accidental death, as reported in a short pamphlet (a sort of dossier on the victims) to be studied twice a day for 7 days (see Fig. 1 for an example).

**Fig. 1:**
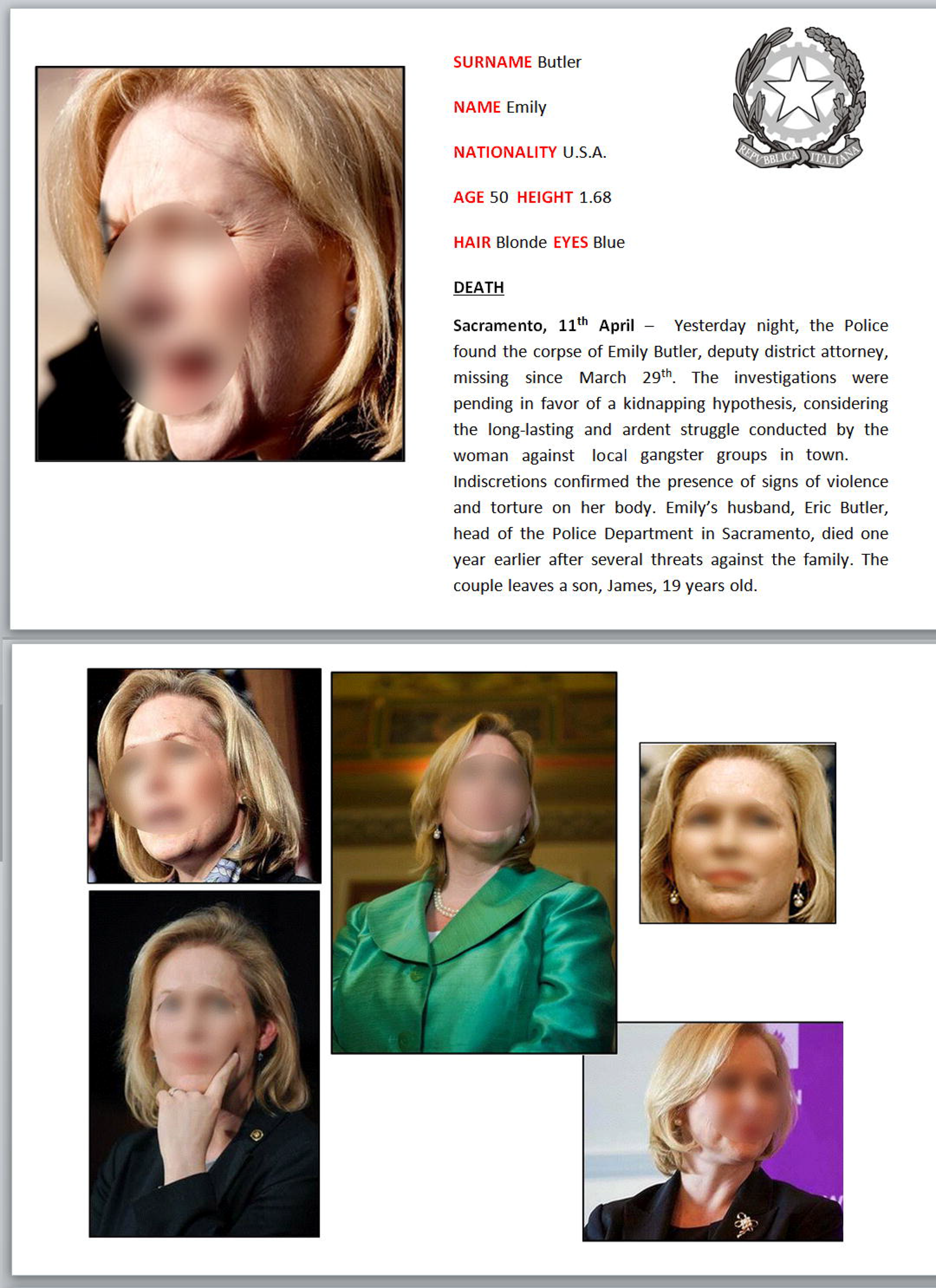
Example of a victim’s dossier to be studied during the encoding phase. Participants were required to study the identikits (victims profile, pictures and description of death circumstances), twice a day for seven days, after waking up and immediately before going bed, for at least 10 minutes. Please note that faces are blurred in the paper (just for publication purposes), but they were unaltered during stimulus presentation.

### Stimuli validation

All characters were rated for familiarity and attractiveness in a validation phase. Validation aimed at proofing that famous faces were easily recognizable and that new and victim faces were absolutely unknown to a sample representative of an Italian population of University students. Similarly, it was ascertained that male and female identities did not differ in terms of familiarity and attractiveness. The whole set consisted of pictures relative to 30 different individuals (30 Caucasian attractive people, of about 45 years old: 10 Old, 10 New and 10 to be studies). The sex was matched across categories (half male, half female, each). Pictures were rated for familiarity and attractiveness by a group of 10 judges, 5 men and 5 women, which were blind to the study’s aim.

Female judges ranged in age from 22 to 25 years (mean age=23.4 years, SD=1.34) and had a high level of education (16.8 years in school, SD=1.1). Males, ranged in age from 22 to 26 years (mean age=24.4 years, SD=1.81), had the same level of education as the females (16.6 years in school, SD=2.19). Judges were asked to rate each face for its familiarity on a 3-point scale (0=not familiar, 1=somewhat familiar, 2=extremely familiar) and for its perceived attractiveness on a 3-point scale (0=not attractive, 1=somewhat attractive, 2=extremely attractive). The pictures were presented for 2 seconds to judges in a randomized order by means of a PowerPoint presentation.

Familiarity and attractiveness ratings were subjected to two separate multifactorial ANOVAs whose factors of variability were 2 within-subject factors: Familiarity, or Attractiveness (Not at all, Somewhat, Extremely) and Sex of faces (Same, Opposite). Sex variable was coded considering subject’s gender with respect to picture’s gender, which could thus be “Same” or “Opposite” sex.

The ANOVA carried out on familiarity scores revealed a significant effect of Familiarity (F2,18=1694, p<0.00001). Post-hoc comparisons indicated how overlearned faces were rated extremely familiar (1.96; SE=0.02) as compared to recent faces (unknown prior to the study) (0.12, SE=0.04; p<0.0001) or new faces (0.05, SE=0.02; p<0.0001) who were rated as absolutely unknown. No effect of face sex was found.

For what concerns perceived attractiveness, no significant difference was found across classes, or as a function of face sex. The mean attractive value of faces was 0.90 (SE=0.08) indicating how, overall, faces were considered fairly attractive (overlearned=0.88, recent=0.88, new=0.91).

For each of the 30 characters 10 different pictures (i.e., head shots in slightly different orientations and hairstyles) were used, for a total of 300 pictures to be shown during the recognition phase (100 new, 100 recent, 100 old). The reason for having 300 different pictures (and not repeating the same 30 pictures) was to avoid short term memory effects such as repetition suppression, habituation, stimulus recognition etc... (Hermann et al., 2016) within the recording session. The 300 final pictures were evaluated for two other possible confounding variables: brightness and facial expression. For what concerns brightness, stimuli’s luminance was measured by means of a Minolta luminance meter, and luminance values were compared across categories by a two-way ANOVA that proved equiluminance between face categories (F2,294=0.43; p=0.65). Photographs had same size (8 cm X 9 cm) and were presented at a visual angle of 4° 1′ 26’’ in length and 4° 31′ 2’’ in height.

For what concerns facial expression, faces were coded by two judges as “neutral” or “smiling”. Then, for each character, the number of photographs including neutral or smiling expressions was assessed: evaluations were then entered into a repeated-measures ANOVA which revealed that the number of neutral and smiling faces were comparable across categories (F2,24=0.12; p=0.89) and face sex (F1,24=3.2; p=0.09).

### Task and Procedure

The task included two different phases: a study phase and a recognition phase. <u>Study phase.</u> During the week preceding the EEG recording participants were required to familiarize with the 10 “victim” characters, twice a day, 5 minutes per session: once in the morning after waking up, and once in the evening before going to bed. They were required to carefully read all the ten dossiers and to try to figure out the person’s characteristics and personality (when alive) and to imagine their death circumstances. The folder contained a description of each fictions character as a written file, one for each person, in form of a police dossier. Each sheet contained on the front a mug shot and a brief description of the person, including personal details like height, age, civil status, family composition, as well as the circumstances and causes of death. On the back, instead, a collection of five further photos, different from those used for the successive recognition phase, was attached to furnish supplementary cues about the person’s appearance and personality (see Fig. 1). Six different pictures of the same characters were used according to the available knowledge that providing multiple photographs of an unfamiliar face improves identity verification accuracy (Menon et al., 2015)

Instructions stated: “*Dear participant, thank you very much for your precious contribution to the study You are asked to carefully read these instructions for the task you are required to perform. In this folder you’ll find the identikits of 10 victims who have died in dramatic and sometimes horrible circumstances. Try to memorize the faces of these characters by studying each dossier, focusing on the photographs. You’ll have to browse the documents twice a day, in the morning and in the evening, for a total of 10 minutes a day. For a good outcome of the experiment a constant and repeated study throughout the entire week is fundamental. We recommend your utmost seriousness in following these instructions. Thanks again for your cooperation”.*

<u>Recognition phase.</u> After the study phase, participants were called to perform a face recognition task associated to EEG/ERPs recording. Participants were comfortably seated in a dark and acoustically/electrically shielded test area in front of a PC computer screen located 114 cm from their eyes. They were instructed to gaze at the center of the screen where a small red circle served as fixation point, and to avoid any eye or body movements during the recording session. The task consisted in an old/new recognition task. Participants were instructed to decide whether they knew the persons whose faces were presented (whether because they were famous stars or because had studied their police dossier) by pressing a response key, as quickly and accurately as possible, with the index finger for answering yes, and with the middle finger for answering no. The two hands were alternated during the recording session and the hand order was counterbalanced across participants. At the beginning of each session, they were told which hand should have been used to respond. The experimental session was preceded by a training session in which faces from the three categories, different from those used for the experimental session, were provided to make sure that the task was correctly understood. The training session included answering with both the two hands, as well. Each face was displayed for 1000 ms in central visual field, with an inter-stimulus interval (ISI) that ranged from 900 to 1100 ms. The EEG was synchronized to the onset of the stimulus.

Overall the experimental session included 6 experimental runs of 60 pictures each (10 pictures for each category: famous male, famous female, male victim, female victim, unknown male, unknown female) and the other 5 by 48 pictures (8 pictures for each category), in a randomized order (see Fig. 2). Behavioural data were recorded by means of EEVoke system (*ANT Software*, Enschede, The Netherlands)

**Fig. 2:**
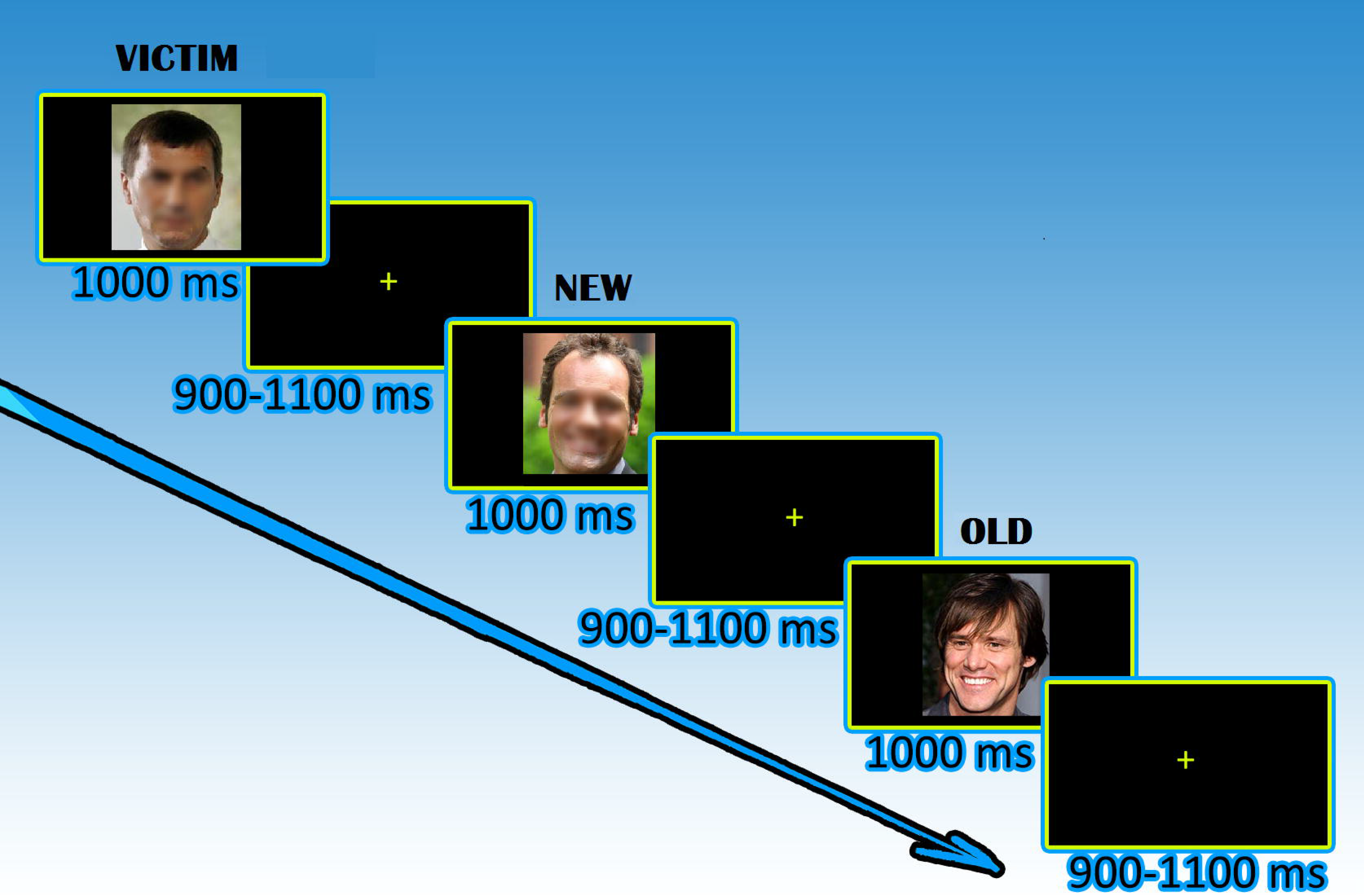
Timeline of stimulus presentation. Each picture was displayed for 1000 ms and followed by a random ISI that ranged from 900 to 1100 ms. The task consisted in deciding whether faces belonged to persons already known, by pressing a response key, as quickly and accurately as possible, with the index finger to signal familiar faces and with the middle finger to signal unknown faces.

### EEG Recording and Analysis

EEG was recorded continuously using the *EEprobe* technology (*ANT Software*, Enschede, The Netherlands) from 128 scalp sites and at the sampling rate of 512 Hz. Horizontal and vertical eye movements were also recorded using the linked mastoids as the reference lead. The EEG and electro-oculogram (EOG) were amplified with a half-amplitude band pass of 0.016−100 Hz. Electrode impedance was maintained below 5 kΩ. EEG epochs were synchronized with the onset of the face presentation. Computerized artifact rejection was performed by means of *EEprobe-ANT* algorithm (Enschede, Netherlands) to discard epochs in which eye movements, blinks, excessive muscle potentials or amplifier blocking occurred. The channels considered for automatic artifact rejection procedure were HEOG, VEOG and FZ. The artifact rejection criterion was a peak-to-peak amplitude that exceeded 50 μV, which resulted in a rejection rate of ~5% (SE=0.65). Evoked-response potentials (ERPs) were averaged from 100 ms before (−100 ms) to 1000 ms after stimulus onset. ERP components were identified and measured with respect to the average baseline voltage over the interval from −100 to 0 ms. Time windows and electrode sites were selected on the basis of where on the scalp and when in time the deflections reached their maximum with respect to the average baseline voltage (following the guidelines for psychophysiological recordings described in in Zani and Proverbio (2003) and Picton et al., (2000). The mean area amplitude of the N200, which reached its maximum amplitude between 200 and 250 ms, and of FN400 component, which reached its maximum amplitude between 350 and 450 ms, were measured at anterior frontal and frontocentral sites (AFF1, AFF2, FC3, FC4). The amplitude of the LPC, which reached its maximum amplitude between 500 and 600 ms, was measured at centroparietal and parietal sites (CP1, CP2, P3, P4). The amplitude of the late anterior frontal negativity (LAN) was quantified between 650 and 850 ms at AF3, AF4, AFp3h, AFp4h electrode sites.

Multifactorial repeated-measures analyses of variance (ANOVA) were applied to ERP amplitude values. Factors were: face familiarity (Overlearned, Recent, New), sex of the face with respect to the viewers (Same, Opposite), electrode (2 levels: AFF1,2 and FC3,4 for N200 and FN400 responses, and CP1,2 and P3,4 for LPC component) and hemisphere (Left, Right) as within factor.

Post-hoc comparisons were performed by using the Tukey test. Reaction times (RTs) that exceeded the mean value ±2 standard deviations were discarded, which resulted in a rejection rate of about 5%. Data normality was assessed through Shapiro-Wilk test (p<0.15).

Accuracy percentages were converted to arcsine values in order to undergo an analyses of variance. As well known (e.g., Snedecor et al., 1989) percentage values do not respect homoscedasticity necessary for ANOVA data distribution and for this reason they need to be transformed in arcsine values. In fact the distribution of percentages is binomial while arcsine transformation of data makes the distribution normal.

The percentage of correct recognitions (hits) for old and victim faces categories, and of correct rejections, for new faces category, were computed. Both RTs and accuracy percentages were subjected to separate multifactorial repeated-measures ANOVAs with 2 within-subject factors: Familiarity (Overlearned, Recent, New) and face Sex (Same, Opposite). The percentage of omissions was negligible (0.87%).

## Results

### Behavioral Results

<u>*Hits*</u> *(familiarity factor)*: The analysis of correct responses (recognition of familiar faces and rejection of unknown faces) revealed a main effect of Familiarity (F2,42=6.06, p<0.005); post-hoc comparisons showed that participants were less accurate in the recognition of new faces (71.17%, SE=2.01) compared to old ones (p<0.005; 78.73%, SE=1.38), while no difference was found in the percentage of correct recognition of overlearned vs. recent faces, new vs. recent faces, or other interaction effects (see Fig. 3).

**Fig. 3:**
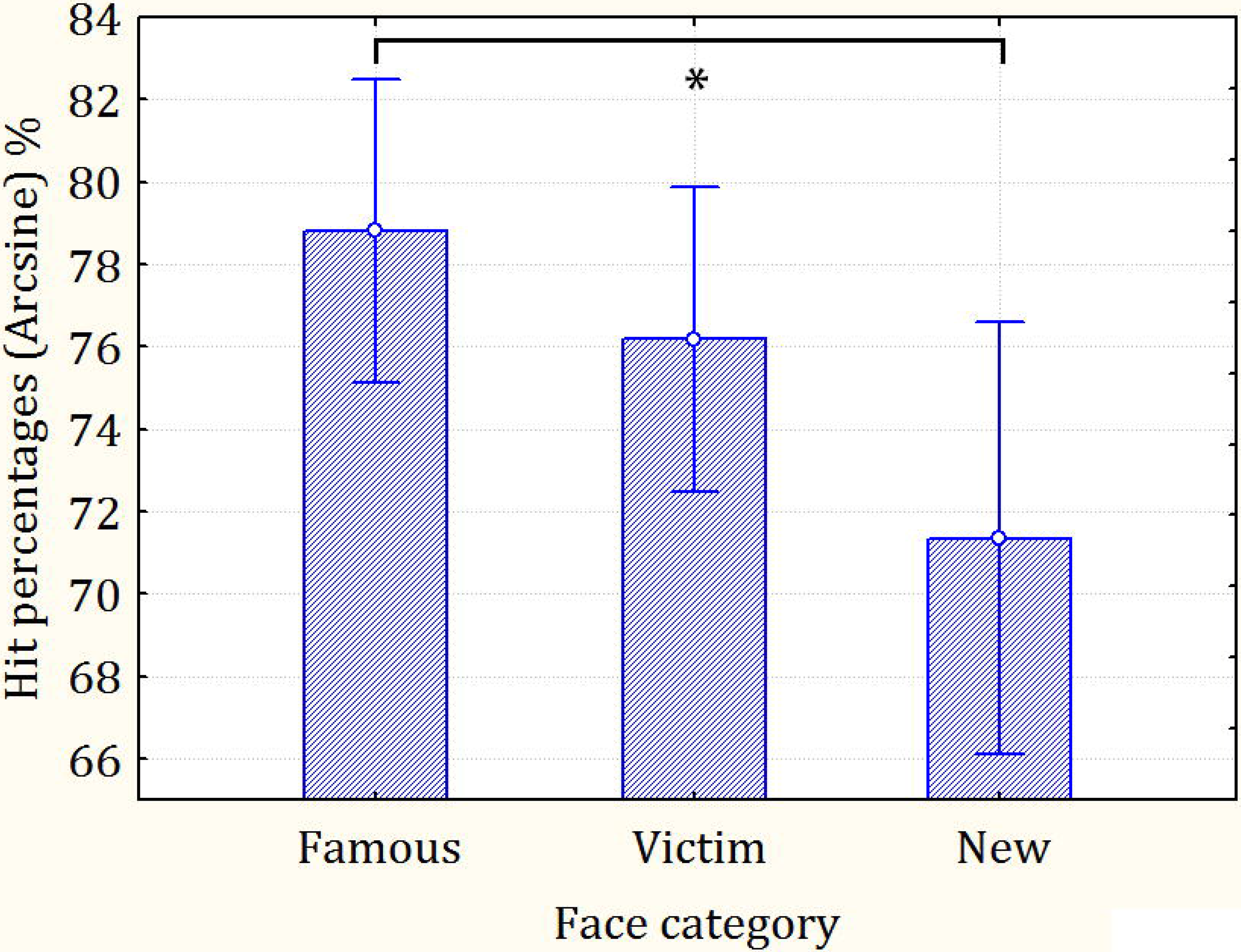
Hit percentages (correct recognitions of overlearned and recent faces and correct rejections of unknown faces).

<u>*RTs*</u> *(familiarity factor):* The ANOVA performed on RTs revealed a significant effect of Familiarity (F2,42=125.53, p<0.00001). Post-hoc comparisons showed that responses were significantly faster (p<0.0001) in response to overlearned (626.22 ms, SE=10.98) than recent faces (668.31 ms, SE=9.79). Furthermore both of them were faster (p<0.0001) than responses to new faces (757.02 ms, SE=14.9).

*(Sex and familiarity x Sex interactions):* Moreover, the analysis revealed a significant effect of Sex (F1,21=5.94, p<0.05) with faster responses for Same-sex (675.54 ms, SE=10.63) compared to Opposite-sex (692.16 ms, SE=12.45) faces. Also, an almost significant interaction effect emerged between Familiarity and Sex (F2,42=3.06, p=0.057); post-hoc comparisons showed that Same-sex faces elicited faster RTs if compared to Opposite-sex ones, but only for recognized faces, whether faces of Victims (Recent: p<0.005, Same: 655.25 ms, SE=9.44; Opposite: M=681.37, SE=11.33) or faces of Famous people (Overlearned: p<0.05; Same: 616.28 ms, SE=11.36; Opposite: 636.16 ms, SE=13.18). Conversely, face sex had no effect when faces belonged to unknown people (p=0.99; Same: 755.08 ms, SE=14.08; Opposite: 758.95 ms, SE=16.44). This effect is clearly visible in Fig. 4.

**Fig. 4:**
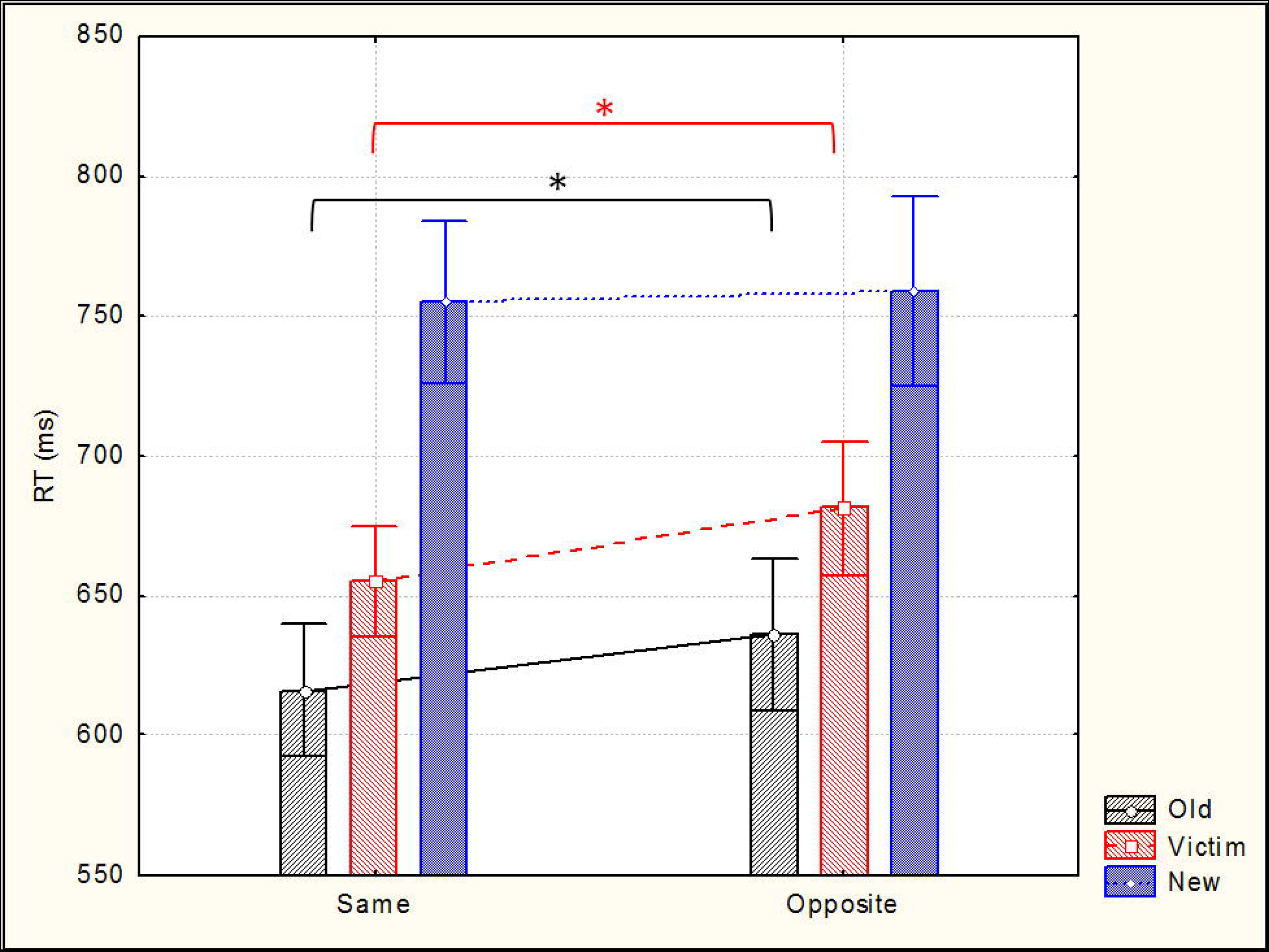
Mean response times (along with standard deviations) as a function of face sex (with respect to that of viewer) and face category.

### Electrophysiological Results

The grand-average ERPs recorded in response to overlearned, recent and new faces are shown in Fig. 5. Responses related to the three kinds of stimuli differed in the amplitude of N200, FN400, LPC and LAN responses, especially over prefrontal sites.

**Fig. 5:**
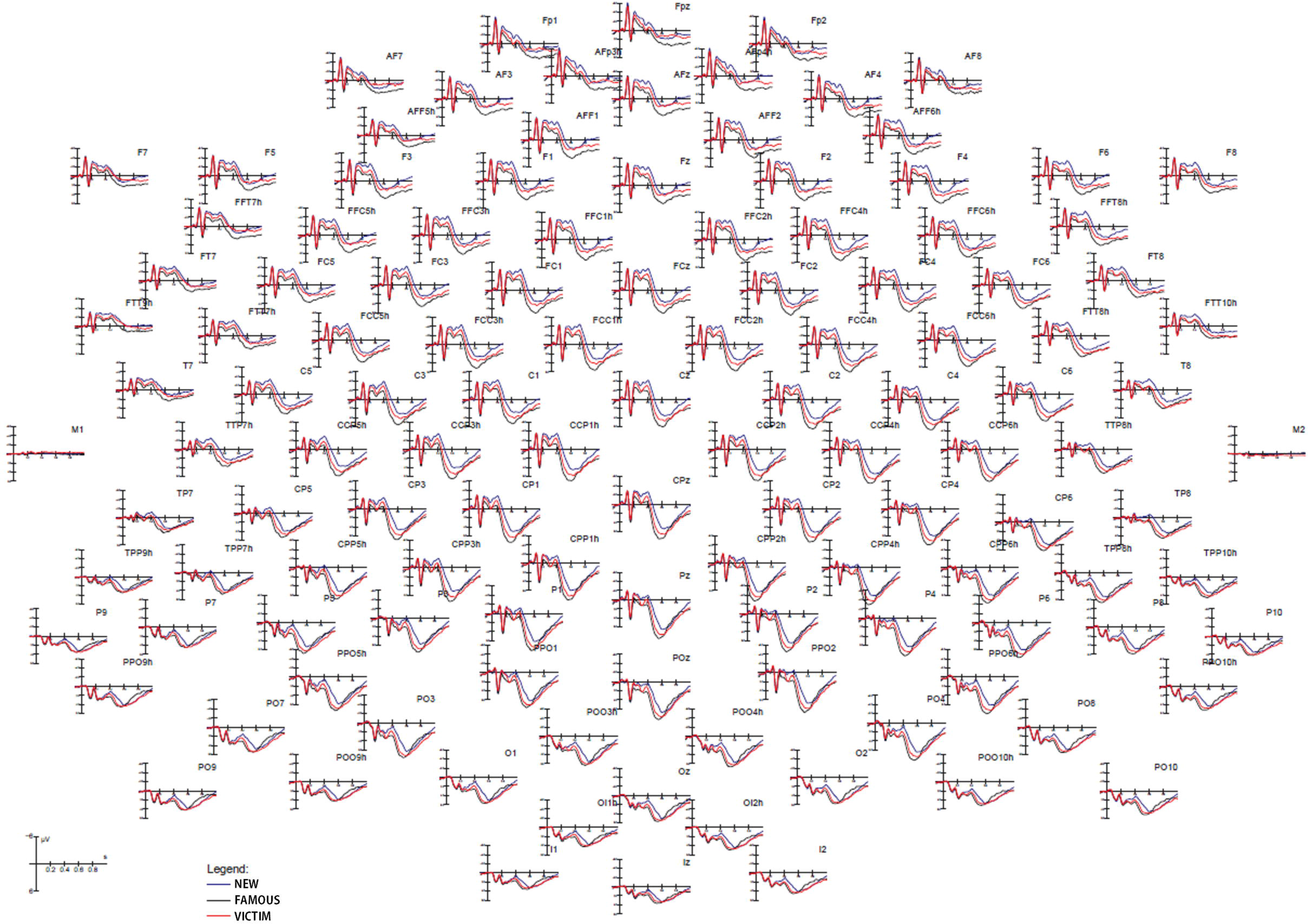
Grand-average ERPs recorded from 128 scalp sites over the left and right hemisphere in response to the 3 face categories.

<u>Anterior N200 response</u>.

(*Familiarity factor*): The ANOVA performed on N200 area amplitudes (measured at anterior frontal and frontocentral sites between 200 and 250 ms) revealed a significant effect of Familiarity (F2,36=8.27, p<0.005) for which processing of new faces was associated with a larger N200 (−4.41 μV, SE=0.93) as compared to overlearned (p<0.001; −3.29 μV, SE=0.87) and recent faces (p<0.001; −3.77 μV, SE=0.87), with no significant difference between the latter. Moreover, the ANOVA revealed a significant effect of electrode (F1,18=8.73, p<0.01) with larger N2 potentials at anterior frontal (−4.15 μV, SE=0.9) than frontocentral (−3.5 μV, SE=0.87) sites.

### FN400 response

*(Familiarity factor):* The ANOVA performed on the FN400 amplitudes (measured at anterior frontal and frontocentral sites between 350 and 450 ms) revealed a significant effect of Familiarity (F2,36=32.66, p<0.00001). Post-hoc analysis showed that the new faces elicited larger FN400 responses (−3.7 μV, SE=0.81) than overlearned (p<0.0005; −1.5 μV, SE=0.78) and recent faces (p<0.0005; −2.13 μV, SE=0.77), while no significant difference was found between the latter. The FN400 modulation can be appreciated in Fig. 6 (upper left).

**Fig. 6:**
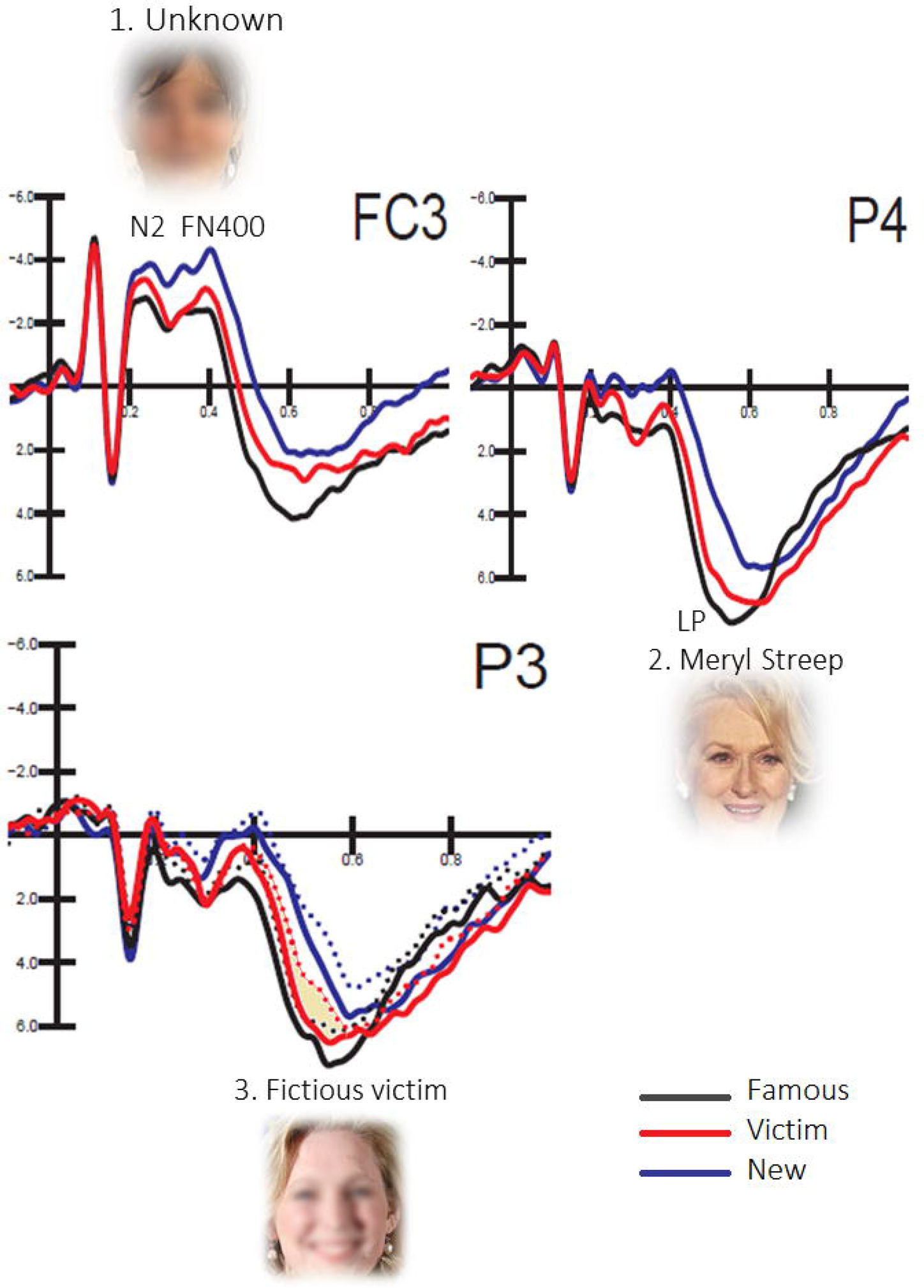
Grand-average ERPs recorded at left frontocentral, left and right parietal sites as a function of face type. Number refers to the functional significance of ERP modulation. Legend: 1. Effect of face Familiarity for N2 and FN400 component. 2. Effect of Temporal recency of memory engram (over right hemispheric areas). 3. Effect of Emotional memory: same-sex effect for victims reveals self-identification and empathic concerns (over left hemispheric areas).

*(Electrode x Hemisphere interaction):* Moreover, a significant interaction effect was found for Electrode x Hemisphere factors (F1,18=6,23, p<0.05): post-hoc comparisons revealed that the largest FN400 response was observed over the left frontocentral sites (all p<0.05; FC3: −2.77 μV, SE=0.77; FC4: −2.1 μV, SE=0.76; AFF1: −2.42 μV, SE=0.82; AFF2: −2.48 μV, SE=0.81), as clearly visible from topographical maps of Fig. 7.

**Fig. 7:**
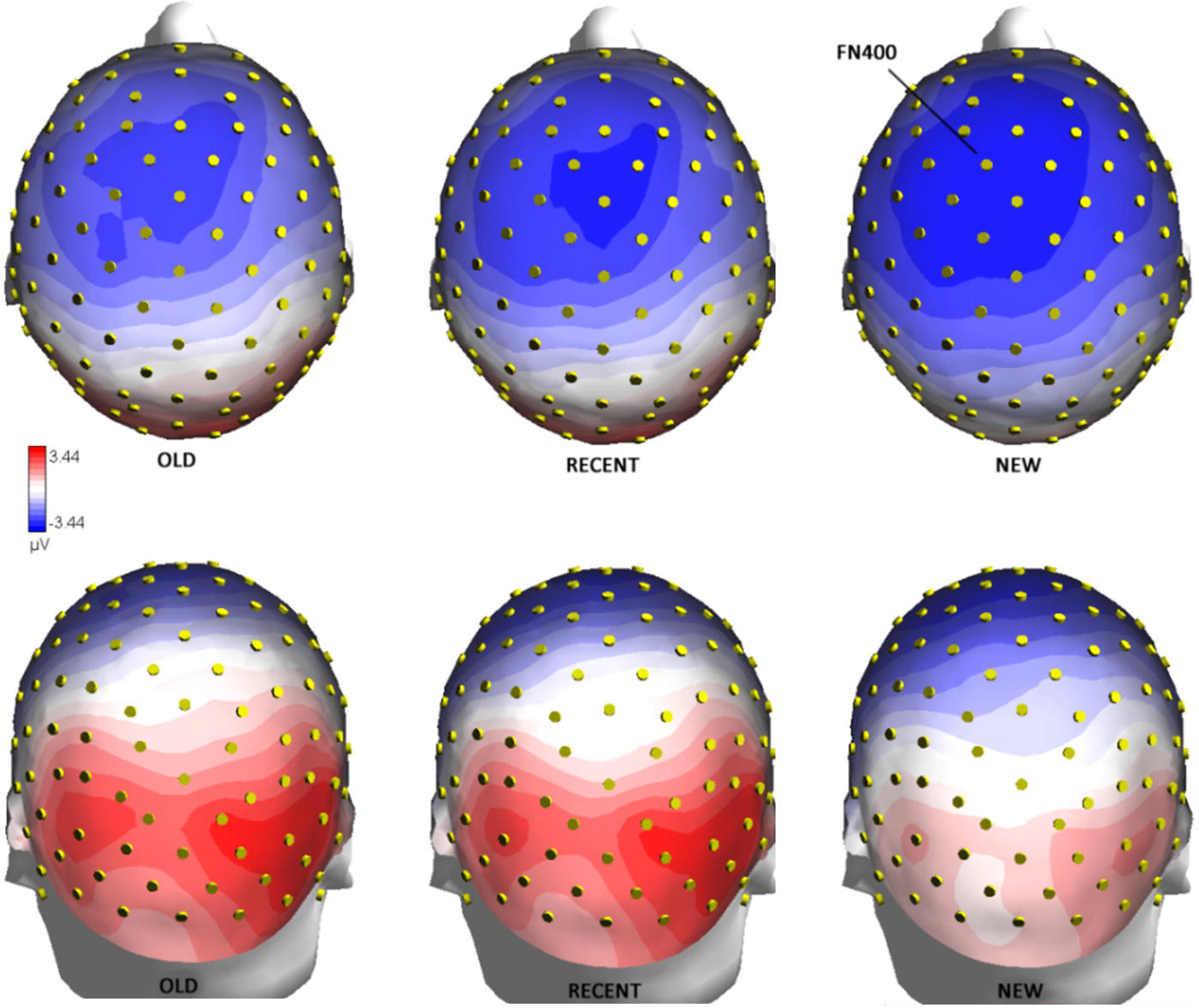
Top (above) and back (below) views of isocolour topographical voltage maps computed in between 350 and 450 ms in response to the 3 stimulus types, during the recognition phase.

*(Familiarity x Sex x Electrode x Hemisphere interaction)*: Finally, post-hoc comparisons of the significant four-way interaction of Familiarity x Sex x Electrode x Hemisphere (F2,36=3.13, p=0.05) showed that Opposite-sex faces elicited larger negativities than Same-sex ones, but only when the faces were familiar. This was true especially for recent faces (Overlearned (p<0.05): Same: −1.3 μV, SE=0.79; Opposite: −1.7 μV, SE=0.84; Recent (p<0.0005): Same: −1.88 μV, SE=0.81; Opposite: −2.39 μV, SE=0.78), while no sex difference was observed for New faces.

### LP response

*(Familiarity factor)*: The ANOVA performed on the LP amplitude (measured at centroparietal and parietal sites between 500 and 600 ms) revealed a significant effect of Familiarity (F2,36=7.43; p<0.00001). In fact overlearned faces (6.56 μV, SE=0.66) elicited larger LP responses compared to recent faces (p<0.005; 5.39 μV, SE=0.54) which, in turn, elicited larger LP potentials than new faces (p<0.0005; 3.15 μV, SE=0.6). LP modulation can be appreciated in Fig. 6 (upper right).

*(Electrode, and Hemisphere factors):* Also, the ANOVA revealed a significant effect of electrode (F1,18=9.32, p<0.01) with larger potentials at parietal (P3/P4: 5.51 μV, SE=0.56) than centroparietal (CP1/CP2: 4.55 μV, SE=0.62) sites. Also significant was the factor Hemisphere (F1,18=5.18, p<0.5), with larger LP responses over the right (5.23 μV, SE=0.58) than the left hemisphere (4.83 μV, SE=0.57).

*(Familiarity x Electrode, and Familiarity x Electrode x Hemisphere interactions):* In addition, significant interactions between Familiarity and Electrode (F2,36=4.29, p<0.05) and Familiarity x Electrode x Hemisphere (F2,36=4.07, p<0.05) were found. Post-hoc comparisons showed that LP to overlearned faces was larger over the right than left parietal area (Overlearned: P4: 7.27 μV, SE=0.68; P3: 6.53 μV, SE=0.63. Recent: P4: 6.18 μV, SE=0.57; P3: 5.59 μV, SE=0.52. New: P4: 3.99 μV, SE=0.58; P3: 3.5 μV, SE=0.59) while the lateralization effect was smaller for other face types and electrodes.

*(Sex factor and interactions)*: Sex factor was also significant (F1,18=8.53, p<0.05) with larger LP in response to same (5.38 μV; SE=0.62) than opposite-sex faces (4.68 μV; SE=0.55). However, the significant four-way interaction of Familiarity x Sex x Electrode x Hemisphere (F2,36=3.34, p<0.05), and relative post-hoc comparisons, revealed that, although, overall, same-sex faces elicited larger LP than opposite-sex faces, the effect was particularly evident for recent faces. The effect emerged at parietal sites where no significant difference in LP amplitude was observed in response to overlearned vs. recent faces if the faces belonged to the same-sex of viewer (P3: p=0.81. Recent-same: 6.15 μV, SE=0.57; overlearned-opposite: 6.31 μV, SE=0.64. P4: p=0.95. Recent-same: 6.72 μV, SE=0.63; overlearned-opposite: 6.86 μV, SE=0.7). The effect of face sex on LP modulation can be appreciated in Fig. 6 (Bottom) and in bargraph of Fig. 8.

**Fig. 8:**
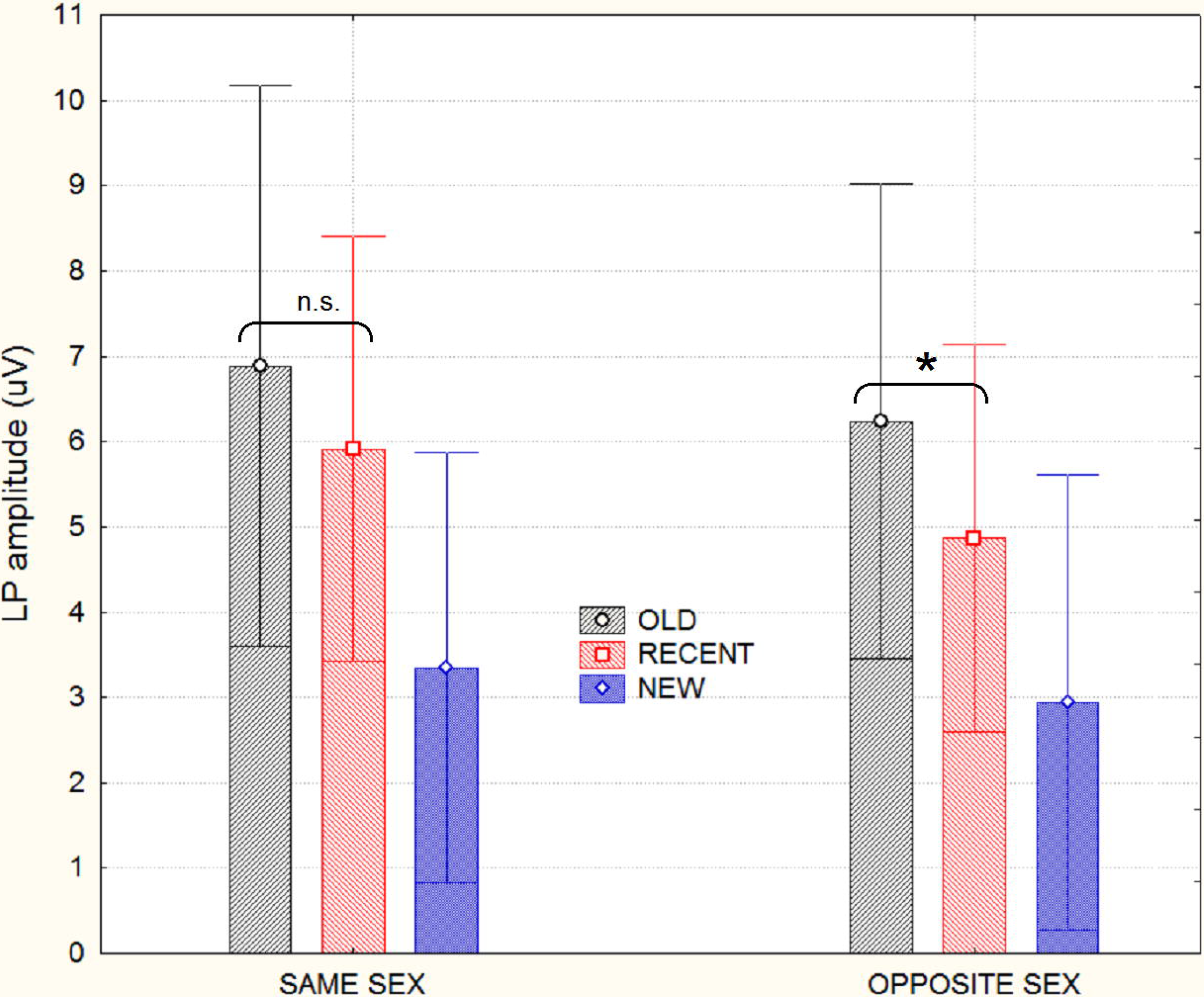
Mean amplitude values (along with standard deviations) of LP response recorded at centroparietal and parietal sites between 500 and 600 ms. Remembered overlearned faces elicited a greater positivity than recent faces of opposite sex and new faces. It can be appreciated how recent faces of same sex elicited an equally larger LP than overlearned faces, thus revealing a selfreferential effect, i.e. an advantage in remembering information more related to self than others.

### Late anterior negativity (650-850 ms)

*(Familiarity factor)*: The ANOVA performed on the LAN amplitudes measured at anterior frontal and prefrontal sites (AF3, AF4, AFp3h, AFp4h) revealed a significant effect of Familiarity (F 2,36=9.2, p<0.00006). In fact overlearned faces (2.77 μV, SE=0.62) elicited much larger positivize responses compared to recent faces (p<0.02; 1.299 μV, SE=0.58) and new faces (p<0.0005; 0.58 μV, SE=0.67), while the difference between the latter did not reach significance. No further effect of electrode, sex or hemisphere was found.

## Discussion

The present paper was aimed at comparing the time course, amplitude and topography of bioelectrical responses elicited during recalling of old and solid information from long term memory (such as the faces of popular movie stars), with that of recently acquired emotional information (faces of fictional characters) as compared to new information (faces of previously unknown people).

Our data showed a modulation of N2 anterior response as a function of familiarity, similarly to FN400 deflection, which agrees with recent memory ERP study (Lawson et al., 2012). It can be assumed that both recently encountered (1 week “old”) and famous faces required less prefrontal processing negativity than new faces at N2 level. Which suggests their similar level of visual familiarity (notwithstanding their profound difference in temporal recency). This interpretation agrees with neuroimaging literature on prefrontal encoding of unfamiliar information (e.g., Lee et al., 2014; Tulving et al., 1996), as well as with electrophysiological literature predicting a smaller anterior negativity to more familiar than new items (e.g., Yonelinas et al., 2010, Proverbio and Orlandi, 2016). Previously, it was assumed that mid latency N2 potential was not particularly sensitive to face familiarity or repetition, but to face structure, being larger in amplitude to faces than scrambled faces or non-face objects (Allison et al., 1999), and, for example, sensitive to face age (being larger to infant than adult faces, Proverbio et al. 2011). However, N200 component has been also related to the degree of face attentional relevance: with N200 amplitudes larger for stimuli with emotional meaning such as faces, more than neutral cues (Streit et al., 2000; Sato et al., 2001). This finding was interpreted as a task-specific emotional marker related to the comprehension of stimulus significance (Herrmann et al., 2002). In the present case, although all stimuli were facial patterns, it could be hypothesized that N2 reflected the different salience of facial information, with novel faces, socially significant stimuli, being coded more deeply than already familiar faces. According to Lawson and coworkers (2112) N2 enhancement would index the detection of stimulus novelty, with more novel and less anticipated items evoking greater N2 amplitudes at frontal sites (e.g., Altenmuller and Gerloff, 1999).

In the next time range FN400 response was also strongly modulated by face familiarity, being larger for new faces than old faces (both overlearned and recent ones). This result is consistent with available literature on face recognition (Curran and Hancock, 2006; Touryan et al., 2011) suggesting that FN400 would reflect stimulus familiarity while later parietal old/new effects (in our study 500-600 ms) would be associated with recollection. The distinction of the two components depends on their distinct topographies on the scalp (Curran, 2000). In fact, in our case, the FN400 reached its maximum amplitude over the frontocentral sites, while the LPC was more evident over parietal sites. However it should mentioned that FN400 might also reflect conceptual priming (deriving from the study of character’s autobiographical information, for recent faces) as argued by some authors (Paller et al., 2007; Wang and Yonelinas, 2012; Jäger et al., 2006, Voss and Paller, 2006).

In our study, accuracy data paralleled FN400 behavior, being sensitive to stimulus familiarity (regardless from recency). Hits percentage to overlearned and recent faces (correct recognitions) were equally greater to those to new faces (correct rejections). On the other hand, response speed data (reflecting the efficiency/certainty of decisional processes), along with LP (parietal late positivity) data, were heavily sensitive to face temporal recency. Indeed, LP amplitude was larger to faces belonging to famous people (overlearned), rather than fictional characters (recent), and both were larger than LP to never encountered faces. In the same vein, RTs were faster to overlearned than recent faces, while unknown faces were processed with a considerable delay. It should be mentioned that 2/3 of the stimuli were old (although with a different degree of recency), and that the “new” vs. “old” response type was therefore not equiprobable, but it is not clear if this might affect performance. Also, it was crucial that overlearned, recent and new faces were actually equiprobable in order for their ERP responses to be comparable. Both RTs and LP measures possible reflected the memory temporal recency, very likely related to engram robustness. Indeed it is possible to assume that face known for many years had stronger and more redundant neural representation, due to the many recollections in different contexts, reactivations and re-coding across time (Kuhl et al., 2012) than faces learned only 7 days before (recent ones). Clinical literature (e.g., Müller et al., 2016) show that in elderly and Alzheimer patients memory loss is reduced for remote compared to more recent life events, and that this also depends on the number of trace reactivations (retrieval frequency of autobiographical memories). LP modulation as a function of face temporal recency recalls the properties of its possible neural generators (see Davachi and DuBrow (2015) for a review), that is the hippocampus and surrounding medial temporal lobe, possibly processing recency and familiarity discrimination necessary for recognition and working memory (Fahy et al., 1993). DuBrow and Davachi (2014) have investigated the role of these areas in representing the sequential order of events in episodic memory. In their fMRI study participants studied sequences of images of celebrity faces and common objects followed by a recency discrimination test. They found that subsequent memory for the order of items in the sequence was related to maintenance of hippocampal BOLD activity patterns across extended sequences. As for animal studies, MacDonald et al. (2011) found specific hippocampal ‘time cells’ that coded distinct moments in time or temporal positions of memory engrams. The temporal firing patterns of these neurons changed when the length of the delay changed. Consistently, Mankin et al. (2012) found a neuronal coding mechanism in the hippocampus of rats that represented the recency of an experience over intervals of hours to days. In their study, when the same event was repeated after such time periods, the activity patterns of hippocampal CA1 cell populations progressively differed with increasing temporal distances. This is of particular interest in the light of the ERP literature indicating the generators of old/new parietal effect (LP) as lying in the hippocampal formation. For example Hoppstädter and coworkers (2015) recorded simultaneous EEG-fMRI signals study during an old/new memory task for words and found that the early frontal effect (350-550 ms) was based on a the dorsolateral prefrontal cortex activation while the late parietal old new effect (580-750ms) was based on the activation of the right posterior hippocampus, parahippocampal cortex and retrosplenial cortex.

In a later range of processing (between 650 and 850 ms) a late frontal negativity revealed a difference between the old items with different sources (overlearned vs recent ones). The scalp distributions of LAN is consistent with the view that the pre-frontal cortex is important for the control of episodic retrieval, and especially source memory (Ranganath and Paller, 1999; Herron and Wilding, 2006; Wilding and Rugg, 1996). Source memory has been specifically studied for example by Wilding and Rugg (1996) in an ERP study in which they compared memory recall of visually presented words in an old/new recognition paradigm. For words recognized as familiar, participants had to determine if they had been spoken by a male or female speaker (i.e. the circumstances of learning, or source memory). ERP results showed that neural activity in the right prefrontal cortex (indexed by an enhanced positivity to old than new words) supported the recovery of contextual information about the item’s study episode.

In our study it might be hypothesized that the much larger positivity (in LAN range) to overlearned than recent faces might relate to their degree of source memory content (since movie stars have been “encountered” repeatedly and in different time periods and circumstances, as opposed to the rather focussed and concentrated 1 week study of recent faces. It should be however mentioned that no overt recall of source memory information was specifically required to perform the present task.

As for the other variables of interest, face sex significantly affected memory recollection especially in the case of autobiographical recent information (“victims”). Effects were found both at behavioral and electrophysiological level. In fact, RTs showed a significant effect of Familiarity x Sex, with faster responses for same- than opposite-sex faces. Previous research on emotions and empathic resonance or mimicry (i.e., an intersubjective induction process by which the observation of an emotion elicits the activation of an analogous representation in the observers (Decety and Meyer, 2008) showed that some specific factors can facilitate emotional mirroring, like perceived similarity; it can influence empathic responding through the tendency to identify more closely with others who appear as being more similar in features such as personality (Gruen and Mendelsohn, 1986) or appearance (Bufalari et al., 2015). This effect was also found, for example, for ingroup versus outgroup identification bias (Tajfel and Turner, 1986) and could have emerged in the case of same-sex faces in the present study. Importantly, same-sex bias was present only in response to familiar persons, both overlearned (p<0.05) but especially recent faces (p<0.005), while RTs to new faces didn’t show a sex effect. It might be assumed that interpersonal identification had a stronger effect in enhancing memory recall of recent than overlearned same sex faces because of a ceiling effect in favor of the latter category, or because identification worked better for ordinary characters depicted as victims (featuring normal jobs such as veterinarian, tourist, writer, entrepreneur, mother of 3 children, University professor, etc..) than movie stars (overlearned faces).

Quite interestingly LP response was larger to familiar faces of viewer’s own sex (either male or female). What is more interesting, nonetheless, is that the effect was particularly evident for recent faces (i.e. victims). Indeed there was no difference in the parietal old/new effect (usually sensitive to the memory trace oldness) recorded in response to overlearned than recent same-sex faces, thus suggesting an effect of affective valence in memory consolidation.

As already discussed in the Introduction, the memory recognition of a face can be influenced by the specific context at encoding time (Megreya & Burton, 2008), the type of the facial expressions (D’Argembeau & Van der Linden, 2011) or the level of the observer empathy (Bate, Parris, Haslam, Kay, 2009). In our case the context provided during encoding was strongly emotional. Observation of victims faces (in 6 different shots) was associated with details relative to their murder, rape or dramatic death. In all cases persons were depicted as very nice and affectionate persons in life, with honorable and estimable professions, leaving small children (e.g. twin baby girls) or inconsolable spouses. Since victim faces were recent, and probably less familiar than movie stars, as known for a shorter time (a week vs. more than 5 years), as also revealed by slower response times and smaller LP amplitudes, the lack of FN400 difference for old vs. recent (victim) faces might be interpreted as an effect of additional emotional information regarding these characters that might (supposedly) reinforced memory encoding.

Interestingly, this effect was more marked in the case of same-sex faces, which could have elicited a higher empathic concern. In fact, as suggested by Westbury and Neumann (2008), perceived similarity can enhance emotional tuning and empathic processes.

More generally, the influence of emotion on memory has been demonstrated by using various types of materials, including complex real-life events (D’Argembeau, Comblain, & Van der Linden, 2005; Talarico, LaBar, & Rubin, 2004), film clips or slide shows (Burke, Heuer, & Reisberg, 1992; Christianson et al.), and lists of words or pictures (Kensinger, 2004; Holland and Kensinger, 2013; LaBar & Cabeza, 2006; Phelps, 2004). It seems that emotionally significant stimuli are processed more efficiently than neutral stimuli. For example, under attentional-blink conditions, emotionally significant words are more often correctly identified than neutral words (Anderson & Phelps, 2001). Also, in brain-damaged patients, visual extinction occurs less often for faces with happy or angry expressions with respect to neutral faces (Vuilleumier & Schwartz, 2001). According to some authors (Richardson et al., 2004) the reason for the enhanced memory of emotional stimuli would depend on an amygdala-based hippocampal modulation. Indeed, while the hippocampus is a critical site for processes related to neuronal encoding, reciprocal modulatory influences between the left amygdala and hippocampus have been demonstrated for effective encoding of emotional material. And indeed amygdala nuclei (in interaction with the hippocampus) would play a crucial role in memory for stimuli that possess affective or motivational significance (Phelps, 2004; Bengner and Malina, 2007).

Overall, these findings hint at crucial role of face emotional valence in increasing the duration of memory retention, without relying on the number of expositions (face popularity), or engram reactivations leading to trace strengthening across months or years, as found in the present study. Indeed, according to neuroscientific literature, memory engrams (i.e., persistent modifications of the synaptic strength via long-term potentiation of pre-existing connections, Poo et al., 2016) are not “imprinted” immediately but evolve over time. Memory strengthening across time would occur by means of repeated and recursive hippocampal sharp wave-ripples responsible of protracted consolidation processes over time.

In conclusion, the present results show that both familiarity and recollection processes contribute to recognition memory, and that these processes are indexed by separate ERP components. However it also shows how emotions and intersubjective identification processes (Decety and Meyer, 2008) toward persons of the same sex are able to modulate memory traces, by increasing the recall of recently acquired items. Indeed, emotional memory in interaction with sex of viewers strengthened the coding of recent information, which resulted in LPs of the same amplitude as to overlearned faces.

## Authors and Contributors

AMP and MEV designed the methods and experiment. MEV and SV prepared stimuli, performed stimuli validation, data acquisition and analyses. AMP interpreted the results and wrote the paper. All authors have contributed to, seen and approved the manuscript.

## Data Accessibility

Anonymized data and details about preprocessing/analyses will be made available to colleagues if requested.

## Acknowledgments

Funded by 9928 013-ATE-0037 grant from University of Milano-Bicocca “How face avoidance affects action understanding and affective processing in Autism”. We are really grateful to Francesco De Benedetto for his technical support.

